# Corticostriatal synaptic weight evolution in a two-alternative forced choice task

**DOI:** 10.1101/549253

**Authors:** C. Vich, K. Dunovan, T. Verstynen, J. Rubin

**Affiliations:** Dept. de Matemàtiques i Informàtica, Universitat de les Illes Balears, Palma, Illes Balears, Spain; Dept. of Psychology, Carnegie Mellon University and Center for the Neural Basis of Cognition, Pittsburgh, PA, USA E-mail: (TV); Dept. of Mathematics and Center for the Neural Basis of Cognition, University of Pittsburgh, Pittsburgh, PA, USA

**Keywords:** Dopamine, reinforcement learning, decision making, spike timing-dependent plasticity

## Abstract

In natural environments, mammals can efficiently select actions based on noisy sensory signals and quickly adapt to unexpected outcomes to better exploit opportunities that arise in the future. Such feedback-based changes in behavior rely on long term plasticity within cortico-basal-ganglia-thalamic networks, driven by dopaminergic modulation of cortical inputs to the direct and indirect pathway neurons of the striatum. While the firing rates of corticostriatal neurons have been shown to adapt across a range of feedback conditions, it remains difficult to directly assess the corticostriatal synaptic weight changes that contribute to these adaptive firing rates. In this work, we simulate a computational model for the evolution of corticostriatal synaptic weights based on a spike timing-dependent plasticity rule driven by dopamine signaling that is induced by outcomes of actions in the context of a two-alternative forced choice task. Results show that plasticity predominantly impacts direct pathway weights, which evolve to drive action selection toward a more-rewarded action in settings with deterministic reward outcomes. After the model is tuned based on such fixed reward scenarios, its performance agrees with the results of behavioral experiments carried out with probabilistic reward paradigms.

## 1 Introduction

The flexible range of mammalian behavior in dynamic and often volatile environments suggests that the neural circuits associated with action selection must be highly modifiable. This adaptive behavior requires that the outcomes of past experiences influence neural circuits in a principled way that maximizes the chances of success in the future [5]. A significant body of experimental work has established that corticostriatal synapses represent one site of such plasticity, which is triggered when a behavior followed by an unexpected reward leads to a change in dopamine levels [8, 39, 44]. Because the corticostriatal synapses represent a key input pathway to the cortico-basal ganglia-thalamic (CBGT) circuits, these dopaminergic changes have been shown to have a critical impact on global network computations related to action selection [2,7,12,15,24,37,51].

In fact, cortical signals to the striatum infiltrate the basal ganglia via two distinct routes, the direct and indirect pathways, each targeted by a corresponding population of striatal medium spiny neurons (MSNs). While these populations are sometimes called D1 (direct) and D2 (indirect) MSNs based on the predominant dopamine receptors that they express, we will refer to these neuron types as dMSNs and iMSNs, respectively. A classic hypothesis posits that the direct pathway provides a go signal that permits an action to be implemented, by disinhibiting downstream targets of inhibitory basal ganglia outputs [33]. Different actions may be driven by dMSNs in different channels, and selection of an action involves cortical activation of the corresponding channel. According to this framework, the indirect pathway promotes inhibitory outputs. When an action associated with one channel is selected, the activity of iMSNs in other channels prevents simultaneous activation of competing actions. According to this theory, if they are active after dMSNs, then the iMSNs in the same channel can terminate previously selected actions [35].

Recent experiments, however, have shown that both the dMSNs and the iMSNs linked with a particular action are simultanteously active during action selection [10, 38, 48, 49]. This co-activation of dMSN and iMSN populations has challenged the traditional model of a strict isomorphism between dMSN activity and excitation and iMSN activity and inhibition. Indeed, more recent theoretical models have proposed that, within an action channel, the dMSN and iMSNs work in a competitive manner to regulate the certainty of a given action decision [3, 14, 15, 31]. For example, Dunovan & Verstynen (2016) proposed a Believer-Skeptic framework for understanding CBGT circuit computations [15]. In this model the direct pathway is cast as the Believer, activated by evidence supporting the favorability of a given action, while the indirect pathway serves as the Skeptic, activated by inputs not in favor of that action. The greater the Believer-Skeptic competition within an action channel, the slower the accumulation of evidence in favor of that action. While this viewpoint shares some similarities with the more classic model [33], it allows for simultaneous increases in dMSN and iMSN populations in a single action channel, corresponding to the accumulation of all types of information relating to that action.

Previous work has developed a computational representation of corticostriatal plasticity in the context of action learning and extinction within the full CBGT circuit [1, 22]. In this framework, corticostriatal synapses are updated based on a spike timing-dependent plasticity (STDP) rule, determined by the timing of a striatal neuron’s spikes relative to the cortical inputs it receives, and on dopamine signals related to reward prediction errors. While the dopamine is shared across neurons and their synapses, the STDP rule sets a synapse-specific eligibility [25, 32, 45], such that only those synapses active with the appropriate timing relative to changes in dopamine are modified.

Here we attempt to adapt and update previous models of dopaminergic learning at the corticostriatal synapses in order to consider how, with repeated evidence presentation, dopamine continuously sculpts synpatic weights at dMSNs and iMSNs in order to influence their relative firing patterns and subsequent behavior. We incorporate dynamically evolving dopamine levels in the setting of either constant or probabilistic rewards delivered in a two-alternative forced choice scenario. In our model, spiking activity of striatal neurons in each action channel is driven by ongoing cortical spike trains (Figure 1), with action selection based on spike patterns at the striatal level resulting from learning-induced weight asymmetries, not from differences in cortical patterns between action channels. Our results predict that the predominant site of corticostriatal plasticity arises at synapses to dMSNs and that emergent striatal activity patterns involve significant spiking in both dMSN and iMSN populations associated with a selected action, consistent with both experimental findings [10, 48, 49] and with competing pathway models like the Believer-Skeptic hypothesis [3, 15, 31].

**Fig. 1.**
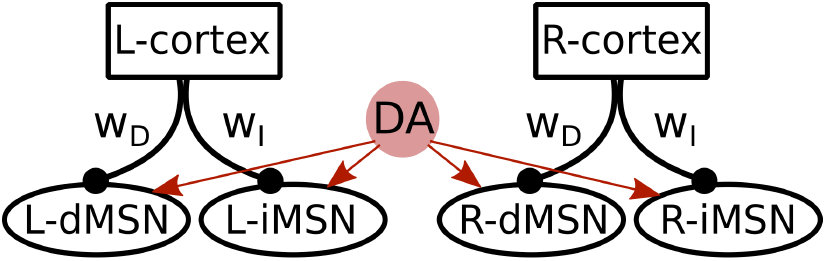
Spike timing-dependent plasticity (STDP) network. The implemented neural network with dopamine (DA) effects on corticostriatal synapses. The *L* and *R* notation denotes the population that influences the choice of the left and right action, respectively. The populations involved are, with *j* ∈ {*L, R*}, *j* – *Cortex*: cortical population, *j* – *dMSN*: direct pathway striatal neurons, and *j* – *iMSN*: indirect pathway striatal neurons. The strengths of these synapses are encoded as weights (*w_j_*) that evolve over time.

The remainder of this paper is organized as follows. In Section 2, we describe our neural model, including cortical spike trains, MSN dynamics, synaptic plasticity, action selection, and reward delivery. We also illustrate how the model effectively functions to update synaptic weights and how it is implemented computationally. In Section 3, we describe model performance when rewards associated with each action are at a fixed level, when rewards are generally fixed but switch after a specific condition is met, and when rewards are probabilistic. The latter scenario allows us to compare our results to experiments with human subjects. Finally, we conclude with a discussion in Section 4 about how our model compares to contemporary theoretical models and emerging empirical observations.

## 2 Methods

The focus of this work is on a network of striatal medium spiny neurons (MSNs) receiving cortical inputs via synapses with plastic weights. In this section we describe the model network (Subsection 2.1) and how the corticostriatal synapses change according to spike timing-dependent plasticity (STDP), which is driven by phasic reward signals resulting from simulated actions and their consequent dopamine release (Subsections 2.2 and 2.3). An example simulation to illustrate the mechanics of the plasticity rule is also presented (Subsection 2.4).

### 2.1 Neural model

We consider a computational model of the striatum consisting of two different populations that receive distinct streams of inputs from the cortex (see Fig. 1, left). Although they do not interact directly, they compete with each other to be the first to select a corresponding action.

Each population contains two different types of units: (i) dMSNs, which facilitate action selection, and (ii) iMSNs, which suppress action selection. Each of these neurons is represented with the exponential integrate-and-fire model, a simplified model that captures the fundamental properties of conductance-based models [19], such that each neural membrane potential obeys the differential equation

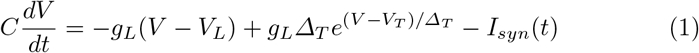

where *g_L_* the leak conductance and *V_L_* the leak reversal potential. In terms of a neural *I* – *V* curve, *V_T_* denotes the voltage that corresponds to the largest input current to which the neuron does not spike in the absence of synaptic input, while Δ_*T*_ stands for the spike slope factor, related to the sharpness of spike initialization. *I_syn_*(*t*) is the synaptic current, given by *I_syn_* (*t*) = *g_syn_* (*t*)(*V*(*t*) – *V_syn_*), where the synaptic conductance *g_syn_*(*t*) changes via a learning procedure (see Subsection 2.2). A reset mechanism is imposed that represents the repolarization of the membrane potential after each spike. Hence, when the neuron reaches a boundary value *V_b_*, the membrane potential is reset to *V_r_*.

The inputs from the cortex to each MSN neuron within a population are generated using a collection of oscillatory Poisson processes with rate *ν* and pairwise correlation c. Each of these cortical spike trains, which we refer to as daughters, is generated from a baseline oscillatory Poisson process {*X*(*t_n_*)}_*n*_, the mother train, which has intensity function λ(1 + *A* sin(2*πθt*)) such that the spike probability at time point *t_n_* is

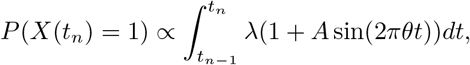

where *A* and *θ* are the amplitude and the frequency of the underlying oscillation, respectively; *t*_*n*+1_ – *t_n_* = : *δt* is the time step; and λ is the mother train rate. After the mother train is computed, each mother spike is transferred to each daughter with probability *p*, checked independently for each daughter. To fix the daughters’ rates and the correlation between the daughter trains, the mother train’s rate is given by *λ* = *ν*/(*p* * *δt*) where

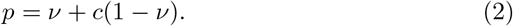

In the STDP network (see Fig. 1, left) for each possible action, we instantiate a corresponding mother train to generate the cortical daughter spike trains for the MSN populations corresponding to that action. Each dMSN neuron or iMSN neuron receives input from a distinct daughter train, with the corresponding transfer probabilities *p^D^* and *p^I^*, respectively. As shown in [28], the cortex to iMSN release probability exceeds that of cortex to dMSN. Hence, we set *p^D^* < *p^I^*.

#### Striatal neuron parameters

We set the exponential integrate-and-fire model parameter values as *C* =1 *μF*/*cm*^2^, *g_L_* = 0.1 *μS/cm*^2^, *V_L_* = −65 *mV*, *V_T_* = —59.9 *mV*, and Δ_*T*_ = 3.48 *mV* (see [19]). The reset parameter values are *V_b_* = –40 *mV* and *V_r_* = –75 *mV*. The synaptic current derives entirely from excitatory inputs from the cortex, so *V_syn_* = 0 *mV*. For these specific parameters, synaptic inputs are required for MSN spiking to occur.

#### Cortical neuron parameters

To compute *p*, we set the daughter Poisson process parameter values as *ν* = 0.002 and *c* = 0.5 and apply equation (2). Once the mother trains are created using these values, we set the iMSN transfer probability to *p^I^* = *p* and the dMSN transfer probability to *p^D^* = 2/3*p^I^*. In most simulations, we set *A* = 0 to consider non-oscillatory cortical activity. We have also tested the learning rule when *A* = 0.06 and *θ* = 25 Hz and obtained similar results.

The network was integrated computationally using the Runge-Kutta (4,5) method in Matlab (ode45) with the time step *δt* = 0.01 *ms*. Different realizations lasting 15 *s* were computed to simulate variability across different subjects in a learning scenario.

Every time that an action is performed (see Subsections 2.3 and 2.4), all populations stop receiving inputs from the cortex until all neurons in the network are in the resting state for at least 50 *ms.* During these silent periods, no MSN spikes occur and hence no new actions are performed (i.e., they are action refractory periods). After these 50*ms*, the network starts receiving synaptic inputs again and we consider a new trial to be underway.

### 2.2 Learning rule

During the learning process, the corticostriatal connections are strengthened or weakened according to previous experiences. In this subsection, we will present equations for a variety of quantities, many of which appear multiple times in the model. Specifically, there are variables *g_syn_,w* for each corticostriatal synapse, *A_PRE_* for each daughter train, *A_POST_* and *E* for each MSN. For all of these, to avoid clutter, we omit subscripts that would indicate explicitly that there are many instances of these variables in the model.

We suppose that the conductance for each corticostriatal synapse onto each MSN neuron, *g_syn_*(*t*), obeys the differential equation

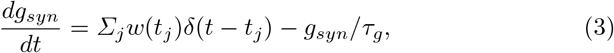

where *t_j_* denotes the time of the *j*th spike in the cortical daughter spike train pre-synaptic to the neuron, *δ*(*t*) is the Dirac delta function, *τ_g_* stands for the decay time constant of the conductance, and *w(t)* is a weight associated with that train at time *t*. The weight is updated by dopamine release and by the neuron’s role in action selection based on a similar formulation to one proposed previously [1], which descends from earlier work [25]. The idea of this plasticity scheme is that an eligibility trace *E* (cf. [45]) represents a neuron’s recent spiking history and hence its eligibility to have its synapses modified, with changes in eligibility following a spike timing-dependent plasticity (STDP) rule that depends on both the pre- and the post-synaptic firing times. Plasticity of corticostriatal synaptic weights depends on this eligibility together with dopamine levels, which in turn depend on the reward consequences that follow neuronal spiking.

To describe the evolution of neuronal eligibility, we first define *A_PRE_* and *A_POST_* to represent a record of pre- and post-synaptic spiking, respectively. Every time that a spike from the corresponding cell occurs, the associated variable increases by a fixed amount, and otherwise, it decays exponentially. That is,

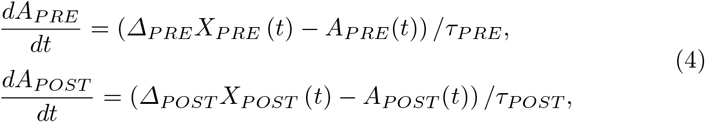

where *X_PRE_*(*t*) and *X_POST_*(*t*) are functions set to 1 at times *t* when, respectively, a neuron that is pre-synaptic to the post-synaptic neuron, or the post-synaptic neuron itself, fires a spike, and are zero otherwise. Δ_*PRE*_ and Δ_*POST*_ are the fixed increments to *A_PRE_* and *A_POST_* due to this firing. The additional parameters τ_*PRE*_,τ_*POST*_ denote the decay time constants for *A_PRE_,A_POST_*, respectively.

The spike time indicators *X_PRE_,X_POST_* and the variables *A_PRE_,A_POST_* are used to implement an STDP-based evolution equation for the eligibility trace, which takes the form

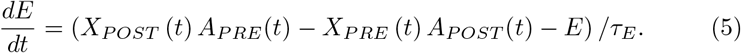

According to this equation, if a pre-synaptic neuron spikes and then its post-synaptic target follows, such that *A_PRE_* > 0 and *X_POST_* becomes 1, then the eligibility *E* increases, while if a post-synaptic spike occurs followed by a pre-synaptic spike, such that *A_POST_* > 0 and *X_PRE_* becomes 1, then *E* decreases. At times without spikes, the eligibility decays exponentially with rate *τ_E_*.

In contrast to some previous work [1], we propose an update scheme for the synaptic weight w(t) that depends on the type of MSN neuron involved in the synapse. It has been observed [13, 21, 26, 40] that dMSNs tend to have less activity than iMSNs at resting states, consistent with our assumption that *p^D^* < *p^I^*, and are more responsive to phasic increases in dopamine than iMSNs. In contrast, iMSNs are largely saturated by tonic dopamine. In both cases, we assume that the eligibility trace modulates the extent to which a synapse can be modified by the dopamine level relative to a tonic baseline (which we without loss of generality take to be 0), consistent with previous models. Hence, we take *w*(*t*) to change according to the equation

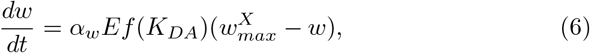

where the function

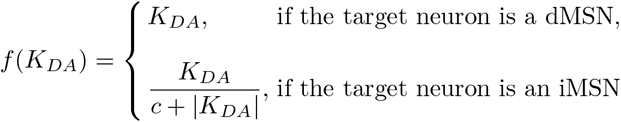

represents sensitivity to phasic dopamine, *α_w_* refers to the learning rate, *K_DA_* denotes the level of dopamine available at the synapses, 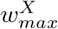 is an upper bound for the weight *w* that depends on whether the postsynaptic neuron is a dMSN (*X* = *D*) or an iMSN (*X* = *I*), *c* controls the saturation of weights to iMSNs, and | ° | denotes the absolute value function. Importantly, we follow past work and take > 0 for dMSNs and *α_w_* < 0 for iMSNs [1]. The form of *f*, chosen to be an odd function for simplicity, may underestimate iMSN sensitivity to decreases in dopamine (e.g., see [31]); because *α_w_* < 0 for iMSNs, this underestimation translates into a weakened increase in weights onto iMSNs, but we shall see that this is not an important factor in our simulation results.

The dopamine level *K_DA_* itself evolves as

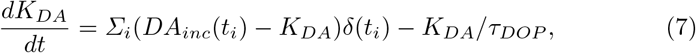

where the sum is taken over the times {*t_i_*} when actions are performed, leading to a change in *K_DA_* that we treat as instantaneous, and *τ_DOP_* is the dopamine decay constant. The DA update value *DA_inc_*(*t_i_*) depends on the performed action as follows:

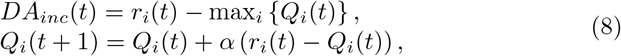

where *r_i_(t)* is the reward associated to action *i* at time *t, Q_i_*(*t*) is an estimate of the value of action *i* at time *t* such that *r_i_*(*t*) – *Q_i_*(*t*) is the subtractive reward prediction error [17], and *α* ∈ [0, 1] is the value learning rate. This rule for action value updates and dopamine release resembles past work [31] but uses a neurally tractable maximization operation (see [27, 41] and references therein) to take into account that reward expectations may be measured relative to optimal past rewards obtained in similar scenarios [9, 34]. In fact, we obtained similar results without the max operation in equation 8, but with slower convergence time (data not shown). The evolution of these variables is illustrated in Fig. 2, which is discussed in more detail in Subsection 2.4.

**Fig. 2.**
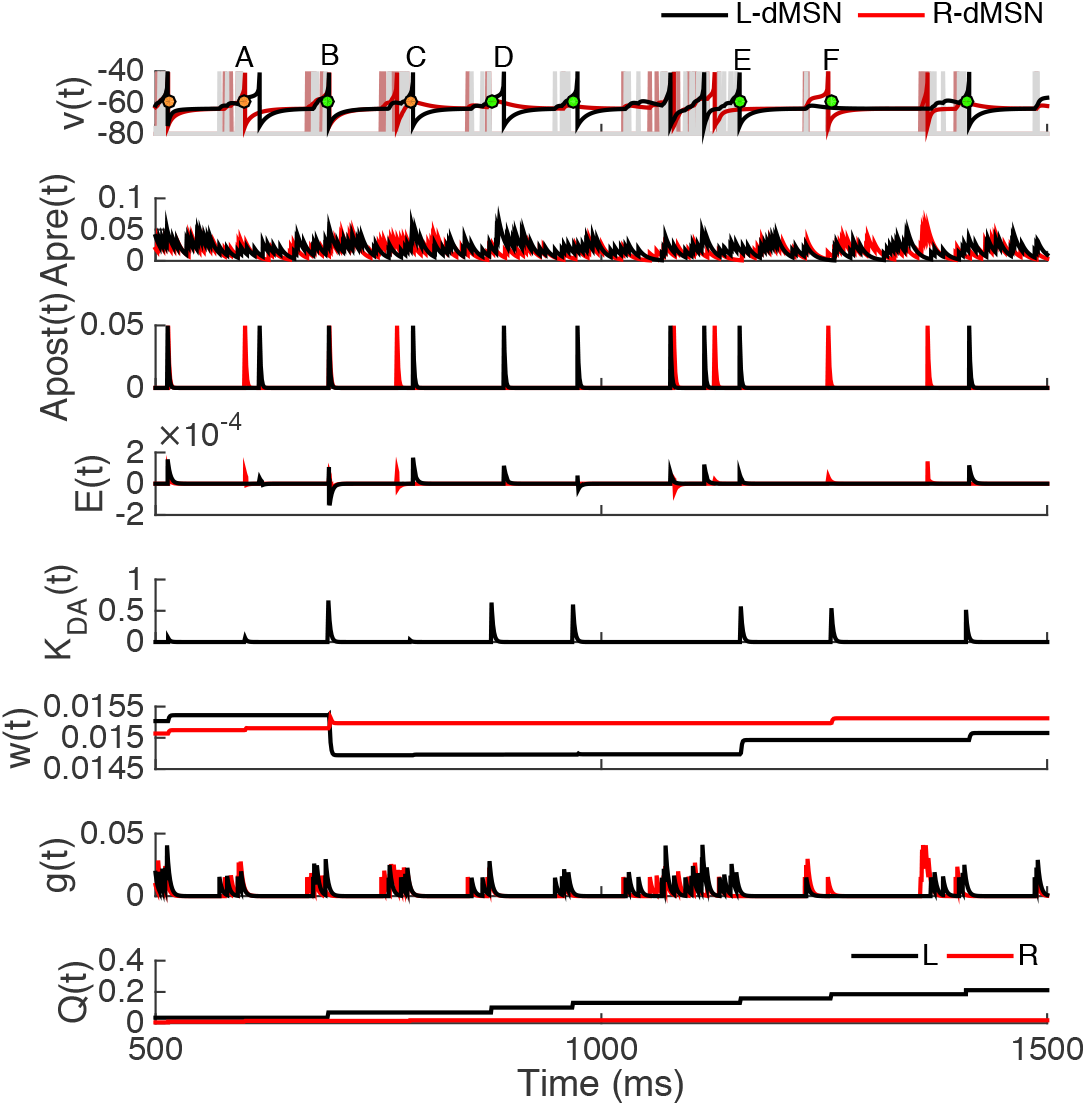
Evolution of the learning rule variables. Learning-related variables were computed for dMSNs in each action channel, one promoting the *L* action (black, actual reward value 0.7) and one promoting the *R* action (red, actual reward value 0.1). Each panel represents corresponding variables for both neurons except *K_DA_*(*t*), which is common across all neurons. For each example neuron, the top panel shows its membrane potential (dark trace) and the cortical spike trains it receives (light trace with many spikes). This panel also represents the action onset times: green and orange dots if actions *L* and *R* occur, respectively. Different example cases labeled with letters (A,B,C,D,E,F) are described in the text in Subsection 2.4.

### 2.3 Actions and rewards

#### Actions

Each dMSN facilitates performance of a specific action. We specify that an action occurs, and so a decision is made by the model, when at least three different dMSNs of the same population spike in a small time window of duration Δ_*DA*_. When this condition occurs, a reward is delivered and the dopamine level is updated correspondingly, impacting all synaptic weights in the network in a way that depends on eligibility as specified in equation (6). Then, the spike counting and the initial window time are reset, and cortical spikes to all neurons are turned off over the next 50 ms before resuming again as usual.

We assume that iMSN activity within a population counters the performance of the action associated with that population [42]. We implement this effect by specifying that when an iMSN in a population fires, the most recent spike fired by a dMSN in that population is suppressed. Note that this rule need not contradict observed activation of both dMSNs and iMSNs preceding a decision [10], see Subsection 3. We also implemented a version of the network in which each iMSN spike cancels the previous spike from both MSN populations. Preliminary simulations of this variant gave similar results to our primary version but with slower convergence (results not shown).

For convenience, we refer to the action implemented by one population of neurons as “left” or *L* and the action selected by the other population as “right” or *R*.

#### Rewards

In our simulations, to test the learning rule, we present results from different reward scenarios. In one case, we use constant rewards, with *r_L_* = 0.7 and *r_R_* = 0.1. In tuning the model, we also considered a regime with reward switches: reward values were as in the constant reward case but after a certain number of actions occurred, the reward-action associations were exchanged. We consider two different switching cases: a solely switch, changing the rewards after 20 *L* actions occurred; and multiple switches, changing the rewards after 15 preferred actions occurred. Finally, we also implemented probabilistic rewards: every time that an action occurs, the reward *r_i_* is set to be 1 with probability *p_i_* or 0 otherwise, *i* ∈ {*L,R*}, with *p_L_* + *p_R_* = 1 and *p_L_* > *p_R_*, keeping the action *L* as the preferred one. Specifically, we consider the three different cases of *p_L_* = 0.85, *p_L_* = 0.75, and *p_L_* = 0.65 to allow comparison with previous results [20].

### 2.4 Example implementation

The algorithm for the learning rule simulations is found in Algorithm 1.

#### Algorithm 1 Dopamine plasticity algorithm

First, generate cortical mother spike trains and extract daughter trains to be used as inputs to each MSN from the mother trains.

Next, while *t* < *t_end_*,

1. use RK45, with step size *dt* = 0.01 *ms*, to compute the voltages of the MSNs in the network at the current time *t* from equations (1) and (3),
2. for each MSN, set the corresponding *X_POST_*(*t*) equal to 1 if a spike is performed or 0 otherwise and set the corresponding *X_PRE_* (*t*) to 1 if an input spike arrives or 0 otherwise,
3. update the *action* condition by checking sequentially for the following two events:

– if any iMSN neuron in population *i* ∈ {*L, R*} spikes, then the most recent spike performed by any of the dMSNs of population *i* is cancelled;
– for each *i* ∈ {*L, R*}, count the number of spikes of the dMSNs in the ith population inside a time window consisting of the last Δ_*DA*_ ms; if at least *n_act_* spikes have occurred in this window, then action i has occurred and we update *DA_inc_* and *Q_i_* according to equation (8),
4. use RK45, with step size *dt* = 0.01 *ms*, to solve equations (4)–(6) for each synapse, along with equation (7) shared by all synapses, yielding an update of DA and all synaptic weight levels; for neurons that have *X_PRE_* (*t*) = 1, update synaptic conductance using *g*(*t*) = *g*(*t*) + *w*(*t*),
5. set *t* = *t* + *dt*.

Fig. 2 illustrates the evolution of all of the learning rule variables over a brief time window. Cortical spikes (thin straight light lines, top panel) can drive voltage spikes of dMSNs (dark curves, top panel), which in turn may or may not contribute to action selection (green – for *L* – and orange – for *R* – dots, top panel). Each time a dMSN fires, its eligibility trace will deviate from baseline according to the STDP rule in equation (5). In this example, the rewards are *r_L_* =0.7 and *r_R_* =0.1, such that every performance of *L* leads to an appreciable surge in *K_DA_*, with an associated rise in *Q_L_*, but performances of *R* do not cause such large increases in *K_DA_* and *Q_R_*.

Various time points are labeled in the top panel of Fig. 2. At time A, *R* is selected. The illustrated *R*-dMSN fires just before this time and hence its eligibility increases. There is a small increase in *K_DA_* leading to a small increase in the *w* for this dMSN. At time B, *L* is selected. Although it is difficult to detect at this resolution, the illustrated *L*-dMSN fires just after the action, such that its *E* becomes negative and the resulting large surge in *K_DA_* causes a sizeable drop in *w_L_*. At time C, *R* is selected again. This time, the *R*-dMSN fired well before time C, so its eligibility is small, and this combines with the small *K_DA_* increase to lead to a negligible increase in *w_R_*. At time D, action L is selected but the firing of the *L*-dMSN is sufficiently late after this that no change in *w_L_* results. At time E, *L* is selected again. This time, the *L*-dMSN fires just before the action leading to a large eligilibity and corresponding increase in *w_L_*. Finally, at time F, *L* is selected. In this instance, the *R*-dMSN fired just before selection and hence is eligible, causing *w_R_* to increase when *K_DA_* goes up. Although this weight change does not reflect correct learning, it is completely reasonable, since the physiological synaptic machinery has no way to know that firing of the *R*-dMSN did not contribute to the selected action *L*.

### 2.5 Learning rule parameters

The learning rule parameters have been chosen to capture various experimental observations, including some differences between dMSNs and iMSNs. First, it has been shown that cortical inputs to dMSNs yield more prolonged responses with more action potentials than what results from cortical inputs to iMSNs [18]. Moreover, dMSNs spike more than iMSNs when both types receive similar cortical inputs [16]. Hence, the effective weights of cortical inputs to dMSNs should be able to become stronger than those to iMSNs, which we encode by selecting 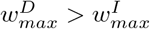. This choice is also consistent with the observation that dMSNs are more sensitive to phasic dopamine than are iMSNs [13, 21, 26, 40]. On the other hand, the baseline firing rates of iMSNs exceed the baseline of dMSNs [30], and hence we take the initial condition for *w*(*t*) for the iMSNs greater than that for the dMSNs.

The relative values of other parameters are largely based on past computational work [1], albeit with different magnitudes to allow shorter simulation times. The learning rate *α_w_* for the dMSNs is chosen to be positive and larger than the absolute value of the negative rate value for the iMSNs. The parameters *Δ_PRE_*, Δ_*POST*_, *τ*_*E*_, *τ_PRE_*, and *τ_POST_* have been assigned the same values for both types of neurons, keeping the relations Δ_*PRE*_ > Δ_*POST*_ and *τ_PRE_* > τ_*POST*_. Finally, the rest of the parameters have been adjusted to give reasonable learning outcomes.

#### Parameter values

We use the following parameter values in all of our simulations: *τ_DOP_* = 2 *ms, Δ_DA_* = 6 *ms, τ_g_* = 3 *ms, α* = 0.05 and *c* = 2.5. For both dMSNs and iMSNs, we set *Δ_PRE_* = 10 (instead of *Δ_PRE_* = 0.1; [1]), *Δ_POST_* = 6 (instead of *Δ_POST_* = 0.006; [1]), *τ_E_* =3 (instead of *τ_E_* = 150; [1]), *τ_PRE_* = 9 (instead of *τ_PRE_* = 150; [1]), and *τ_POST_* = 1.2 (instead of *τ_POST_* = 3; [1]). Finally, *α_w_* = {80, −55} (instead of *α_w_* = {12, −11}; [1]) and *w_max_* = {0.1, 0.03} (instead of *w_max_* = {0.00045, 0}; [1]), where the first value refers to dMSNs and the second to iMSNs. Note that different reward values, *r_i_*, were used in different types of simulations, as explained in the associated text.

#### Learning rule initial conditions

The initial conditions used to numerically integrate the system are *w* = 0.015 for weights of synapses to dMSNs and *w* = 0.018 for iMSNs, with the rest of the variables relating to value estimation and dopamine modulation initialized to 0.

### 2.6 Definitions of quantities computed from the STDP model

#### Averaged population firing rate

We compute the firing rate of a neuron by adding up the number of spikes the neuron fires within a time window and dividing by the duration of that window. The averaged population firing rate is compute as the average of all neurons’ firing rates over a population, given by

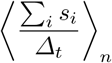

where Δ_*t*_ is the time window in *ms, s_i_* is the spike train corresponding to neuron *i*, and 〈·〉_*n*_ denotes the mean over the n neurons in the population, The time course of the population firing rate is computed this way, using a disjoint sequence of time windows with *Δ_t_* = 500 *ms*.

#### Action frequency

We compute the rate of a specific action i in a small window of *Δ* = 500 *ms* as the number of occurrences of action i within that window divided by *Δ*.

#### Mean behavioral learning curves across subjects

The behavioral learning curves indicate, as functions of trial number, the fraction of trials on which the more highly rewarded action is selected. Within a realization, using a sliding trial count window of 5 trials, we computed fraction of preferred actions selected (number of preferred actions divided by the total number of actions). Then we averaged over *N* realizations.

#### Evolution of the mean (across subjects) difference in model-estimated action values

Using *N* different realizations (simulating subjects in a behavioral experiment), we computed the difference of the expected reward of action *L* and the expected reward of action *R* at the time of each action selection (that is, *Q_L_*(*t*\*) − *Q_R_*(*t*\*), where *t*\* is the time of action selection). Notice that *Q_i_*(*t*\*), for *i* ∈ {*L, R*}, only changes when an action occurs. Moreover, to average across realizations, we only considered the action number rather than the action onset time.

## 3 Results

To evaluate how dopaminergic plasticity impacts the efficacy of corticostriatal synapses, we modeled learning using a spike timing-dependent plasticity (STDP) paradigm in a simulation of corticostriatal networks implementing a simple two-alternative forced choice task. In this scenario, one of two available actions, which we call left (*L*) and right (*R*), was selected by the spiking of model striatal medium spiny neurons (MSNs; Subsection 2.3). These model MSNs were grouped into action channels receiving inputs from distinct cortical sources (Fig. 1, left). Every time an action was selected, dopamine was released, after a short delay, at an intensity proportional to a reward prediction error [equations (7) and (8)]. All neurons in the network experienced this non-targeted increase in dopamine, emulating striatal release of dopamine by substantia nigra pars compacta neurons, leading to plasticity of corticostriatal synapses [equation (6); see Fig. 2].

The model network was initialized so that it did not *a priori* distinguish between *L* and *R* actions. We first performed simulations in which a fixed reward level was associated with each action, with *r_L_* > *r_R_*, to assist in parameter tuning and verify effective model operation. Next, we continued with the constant reward scenario but with reward values exchanged (i.e., *L* becomes the non-preferred action while *R* becomes the most rewarded one) after a certain number of actions, to see if the network is capable of learning that the reward switch has occurred and representing how long it lasts. We finally turn to results obtained with probabilistic rewards, as described in the last paragraph in Subsection 2.3, to compare with data from experiments with human subjects.

### 3.1 Constant rewards scenario

In this first scenario, where *r_L_* = 0.7 and *r_R_* = 0.1, a gradual change in corticostriatal synaptic weights occurred (Fig. 3A) in parallel with the learning of the actions’ values (Fig. 3B).

**Fig. 3.**
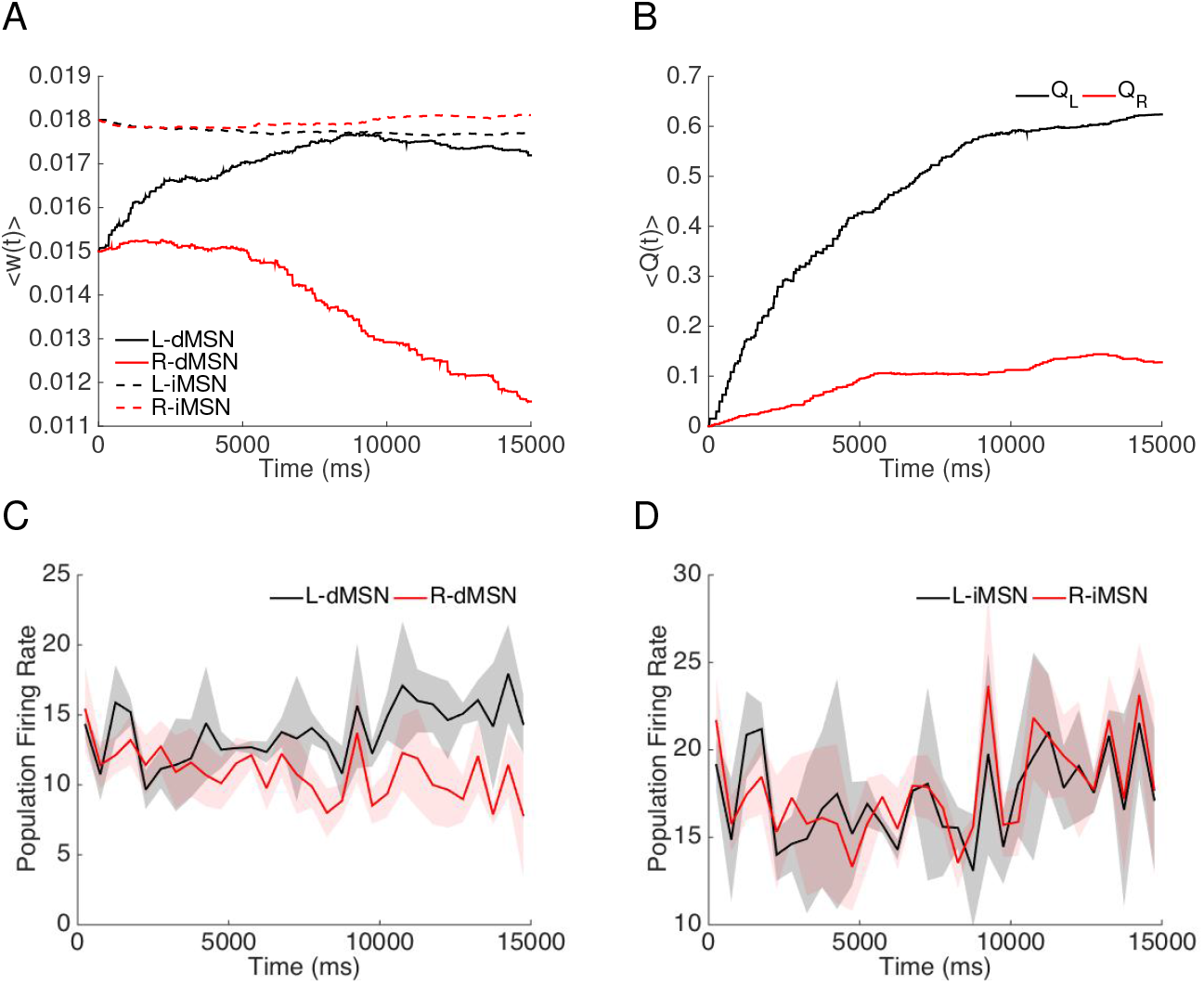
STDP simulations with constant reward feedback. Time courses of corticostriatal synapse weights and firing rates are shown for simulations performed with a constant (i.e., non-probabilistic) reward schedule, where (*r_L_*(*t*) = 0.7 and *r_R_*(*t*) = 0.1). A: Averaged weights over 7 different realizations and over each of the four specific populations of neurons, which are dMSN selecting action *L* (solid black); dMSN selecting action *R* (solid red); iMSN countering action *L* (dashed black); iMSN countering action *R* (dashed red). B: Averaged evolution of the action values *Q_L_* (black trace) and *Q_R_* (red trace) over 7 different realizations. C: Firing rates of the dMSN populations selecting actions *L* (black) and *R* (red) over time. D: Firing rates of the iMSN populations countering actions *L* (black) and *R* (red) over time. Data in C,D was discretized into 50 *ms* bins. The transparent regions depict standard deviations.

These changes in synaptic weights induced altered general MSN firing rates (Fig. 3C,D), reflecting changes in the sensitivity of the MSNs to cortical inputs in a way that allowed the network to learn over time to select the more highly rewarded action (Fig. 4A). That is, firing rates in the dMSNs associated with the more highly rewarded action increased, leading to a more frequent selection of that action. On the other hand, firing rates of the iMSNs remained quite similar (Fig. 3C,D). This similarity is consistent with recent experimental results [11], while the finding that dMSNs and iMSNs associated with a selected action are both active has also been reported in several experimental works [10,48,49].

**Fig. 4.**
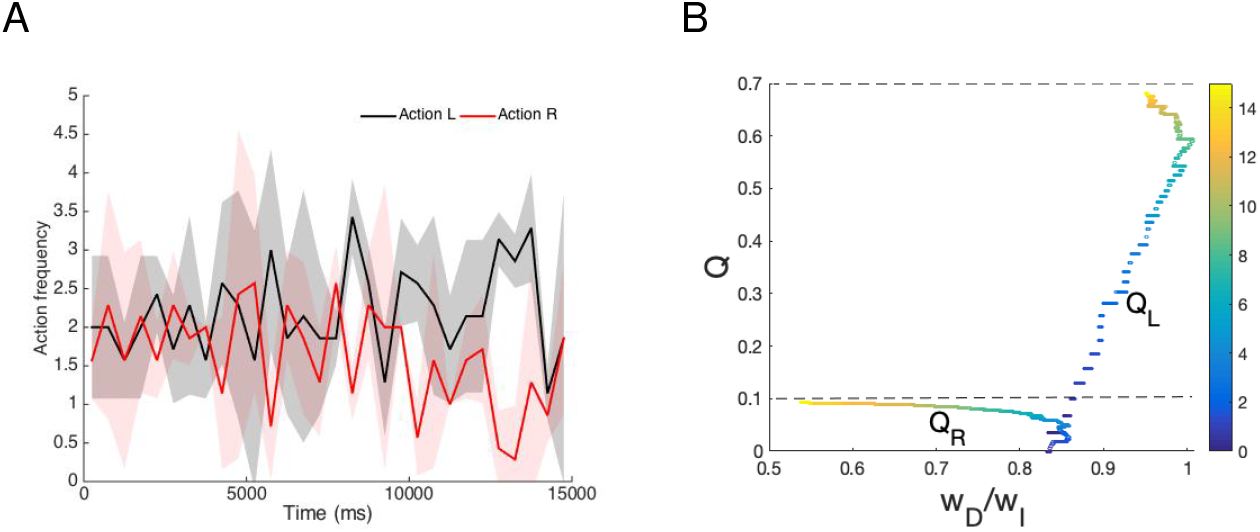
Constant reward task. A: Frequency of performance of *L* (black) and *R* (red) actions is plotted over time (discretized each 50 *ms*) when the rewards are held constant (*r_L_* = 0.7, *r_R_* = 0.1). Both traces are averaged across 7 different realizations. The transparent regions depict standard deviations. B: Estimates of the values of *L* (*Q_L_*) and *R* (*Q_R_*) versus the ratio of the corticostriatal weights to those dMSNs that facilitate the action and those iMSNs that interfere with the action. Each trajectory is colored to show the progression of time. Even without full convergence of the action values *Q_R_* and *Q_L_* to their respective actual reward levels (B), a clear separation of action selection rates emerges (A).

In this model, indirect pathway activity counters action selection by cancelling direct pathway spiking (Subsection 2.3). Based on this cancellation, the ratio of direct pathway weights to indirect pathway weights provides a reasonable representation of the extent to which each action is favored or disfavored.

In our simulations, after a long period of gradual evolution of weights and action values, the direct pathway versus indirect pathway weight ratio of the channel for the less favored action started to drop more rapidly, indicating the emergence of certainty about action values and a clearer separation between frequencies with which the two actions were selected (Fig. 4).

### 3.2 Reward switching scenario

To test whether the network remains flexible after learning a specific action-value relation, we ran additional simulations using a variety of reward schedules in which the reward values associated with the two actions were swapped after the performance of a certain number of actions.

We first performed a simulation in which the rewards associated with the *L* and *R* actions were switched only one time after 5 *s*. In Fig. 5, we can see that when the L-action is rewarded (up to time *t* ≈ 5*s*), the firing rate, action frequency and the action values *Q*(*t*) for the L-dMSNs become higher than those for the *R*-dMSNs, showing a learning of the *L* action. Up to time 5 *s*, 20 *L* actions have been performed and the learning is almost consolidated, since *Q_L_*(*t*) and *Q_R_*(*t*) are close to the actual reward values, *r_L_* = 0.7 and *r_L_* = 0.1, respectively (see top panel of Fig. 5).

**Fig. 5.**
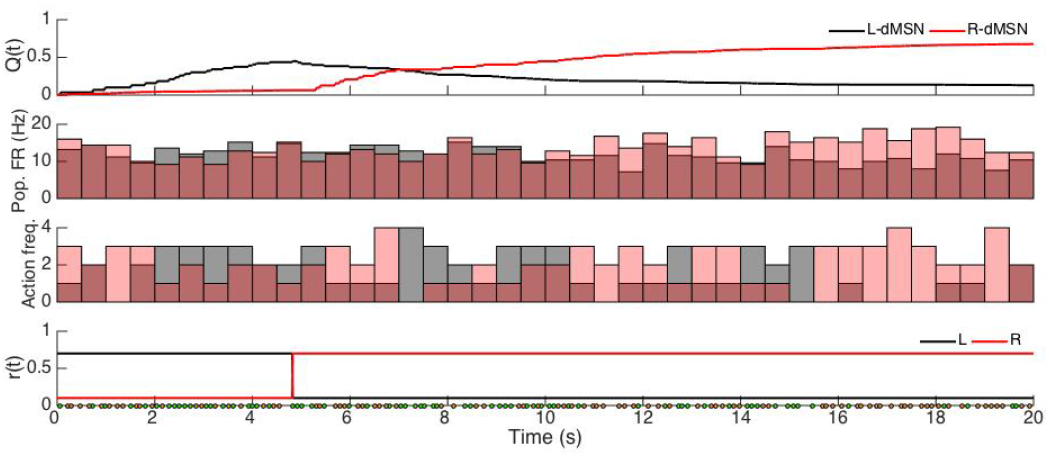
STDP effects following change in action-value associations. The STDP model was simulated on a task in which the associated rewards for *L* and *R* actions are flipped after an initial learning period. The first three panels represent, from top to bottom, the action values (*Q*(*t*)), the firing rates of dMSN neurons for each action (*L*, black; *R*, red), and the action frequency for the dMSN population of neurons that produces the *L* action (black) and the *R* action (red). The bottom panel represents the actual reward values for *L* (black) and *R* (red). The reward values switch when 20 *L* actions have occurred.

At this time, the reward values are swapped. Afterwards, the network is able to learn that the *R* action elicits the preferable reward. Specifically, as we can see in the top panel of Fig. 5, *Q_L_* and *Q_R_* begin evolving toward the new reward levels, switching their relative magnitudes relatively quickly (i.e., in less than 3s) along the way. Although the weights of corticostriatal synapses to *L*-dMSNs (*R*-dMSNs) correspondingly weaken (strengthen), it takes longer, at least 5*s*, until the *R* action is reliably performed more frequently than the *L* action. Thus, the network is able to overcome previously learned contingencies and adaptively learn new ones, yet there is a delay relative to the learning that occurs without the previous bias.

On the other hand, given that the network is capable of learning new optimal actions after the switch, we also wanted to see what happens if the rewards are swapped back and forth before the new learning is consolidates. In Fig. 6, we plot the results of a simulation where the reward values are switched each time that 15 preferred actions take place.

**Fig. 6.**
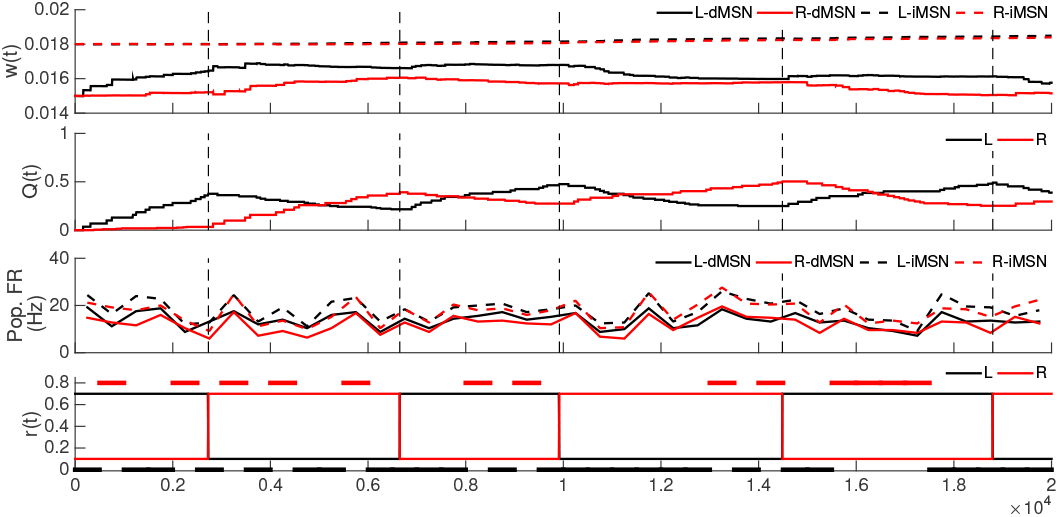
STDP effects after repeatedly switching the highest rewarded action. The STDP model was simulated on a task in which the associated rewards for *L* and *R* actions are flipped each time that 15 preferred actions have occurred. The first three panels represent, from top to bottom, the weights (*w*(*t*)) for the different populations averaged over the number of neurons in each population, the estimates of the action values (*Q*(*t*)), and the firing rates for the different populations averaged over the neurons in each population. The bottom panel represents the actual reward values (thin curves) and the time intervals where an specific action has higher frequency (thick curves). In all panels, black traces refer to the left (*L*) action while red traces indicate the right (*R*) action. In the weight and firing rate plots, the solid lines refer the dMSN neurons while dashed lines refer to iMSN neurons. Vertical dashed lines indicate the times when reward values are switched.

In Fig. 6, after each switch, the estimated action value *Q*(*t*) for the (now) non-preferred action starts to gradually fall off, causing small decreases in the weights of synapses to dMSNs associated with that action and in the mean firing rate of the dMSNs. At the same time, the weights of synapses onto the dMSNs that allow the (now) preferred action and their firing rates increase. In contrast to the action value estimates, which switch quickly, the STDP rule yields a delay in switching of relative synaptic weight values onto dMSNs (top panel of Fig. 6), such that the reward changes are not clearly encoded in the action selection outcomes (bottom panel of Fig. 6). As weights come closer before the next switch, action selection rates equalize. These results illustrate the lag in the STDP rule, which is advantageous for avoiding changes in action policy due to occasional spurious outcomes but requires repeated exposure to learn new reward contingencies, and suggest that some other plasticity mechanism is likely involved in more rapid or one-shot learning.

### 3.3 Probabilistic rewards scenario

While our previous simulations show that applying a dopaminergic plasticity rule to corticostriatal synapses allows for a simple network to learn action values linked to reward magnitude, many reinforcement learning tasks rely on estimating reward probability (e.g., two-armed bandit tasks). To evaluate the network’s capacity to learn from probabilistic rewards, we simulated a variant of a probabilistic reward task and compared the network performance to previous experimental results on action selection with probabilistic rewards in human subjects [20]. For consistency with experiments, we always used *p_L_* + *p_R_* = 1, where *p_L_* and *p_R_* were the probabilities of delivery of a reward of size *r_i_* = 1 when actions *L* and *R* were performed, respectively. Moreover, as in the earlier work, we considered the three cases *p_L_* = 0.65 (high conflict), *p_L_* = 0.75 (medium conflict) and *p_L_* = 0.85 (low conflict).

As in the constant reward case, the corticostriatal synaptic weights onto the two dMSN populations clearly separated out over time (Fig. 7). The separation emerged earlier and became more drastic as the conflict between the rewards associated with the two actions diminished, i.e., as reward probabilities became less similar. Interestingly, for relatively high conflict, corresponding to relatively low *p_L_*, the weights to both dMSN populations rose initially before those onto the less rewarded population eventually diminished. This initial increase likely arises because both actions yielded a reward of 1, leading to a significant dopamine increase, on at least some trials. The weights onto the two iMSN populations remained much more similar. One general trend was that the weights onto the *L*-iMSN neurons decreased, contributing to the bias toward action *L* over action *R*.

**Fig. 7.**
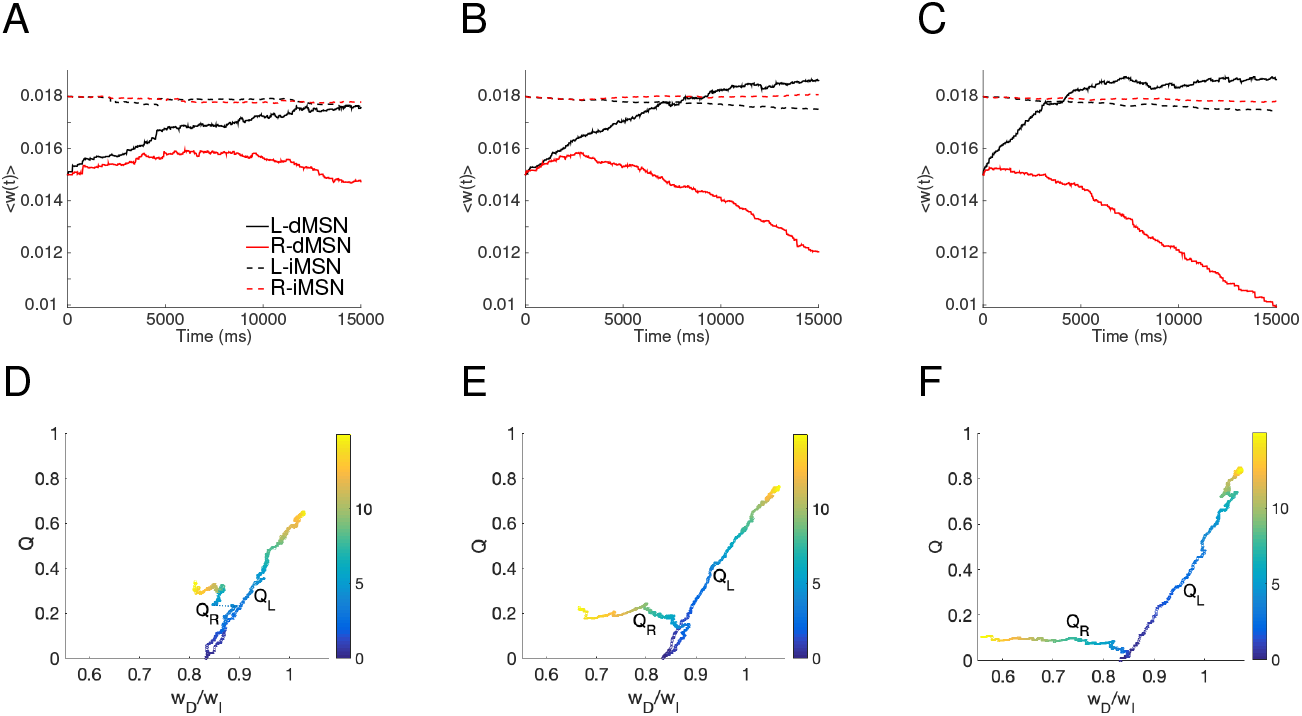
Corticostriatal synaptic weights with probabilistic reward feedback. First column: *p_L_* = 0.65; second column: *p_L_* = 0.75; third column: *p_L_* = 0.85 case. A, B, and C: Averaged weights over each of four specific populations of neurons, which are dMSN neurons selecting action *L* (solid black); dMSN neurons selecting action *R* (solid red); iMSN neurons countering action *L* (dashed black); iMSN neurons countering action *R* (dashed red). D, E, and F: Evolution of the estimates of the value *L* (*Q_L_*) and *R* (*Q_R_*) versus the ratio of the corticostriatal weights to those dMSN neurons that facilitate the action versus the weights to those iMSN that interfere with the action. Both the weights and the ratios have been averaged over 8 different realizations. The jump in *Q_R_* for *p_L_* = 0.65, joined by a dashed line, comes from the time discretization and averaging.

In all three cases, the distinction in synaptic weights translated into differences across the dMSNs’ firing rates (Fig. 8, first row), with *L*-dMSN firing rates (*D_L_*) increasing over time and *R*-dMSN firing rates (*D_R_*) decreasing, resulting in a greater difference that emerged earlier when *p_L_* was larger and hence the conflict between rewards was weaker. Notice that the *D_L_* firing rate reached almost the same value for all three probabilities. In contrast, the *D_R_* firing rate tended to decrease more over time as the conflict level decreased. As expected, based on the changes in corticostriatal synaptic weights, the iMSN population firing rates remained similar for both action channels, although the rates were slightly lower for the population corresponding to the action that was more likely to yield a reward (Fig. 8F).

**Fig. 8.**
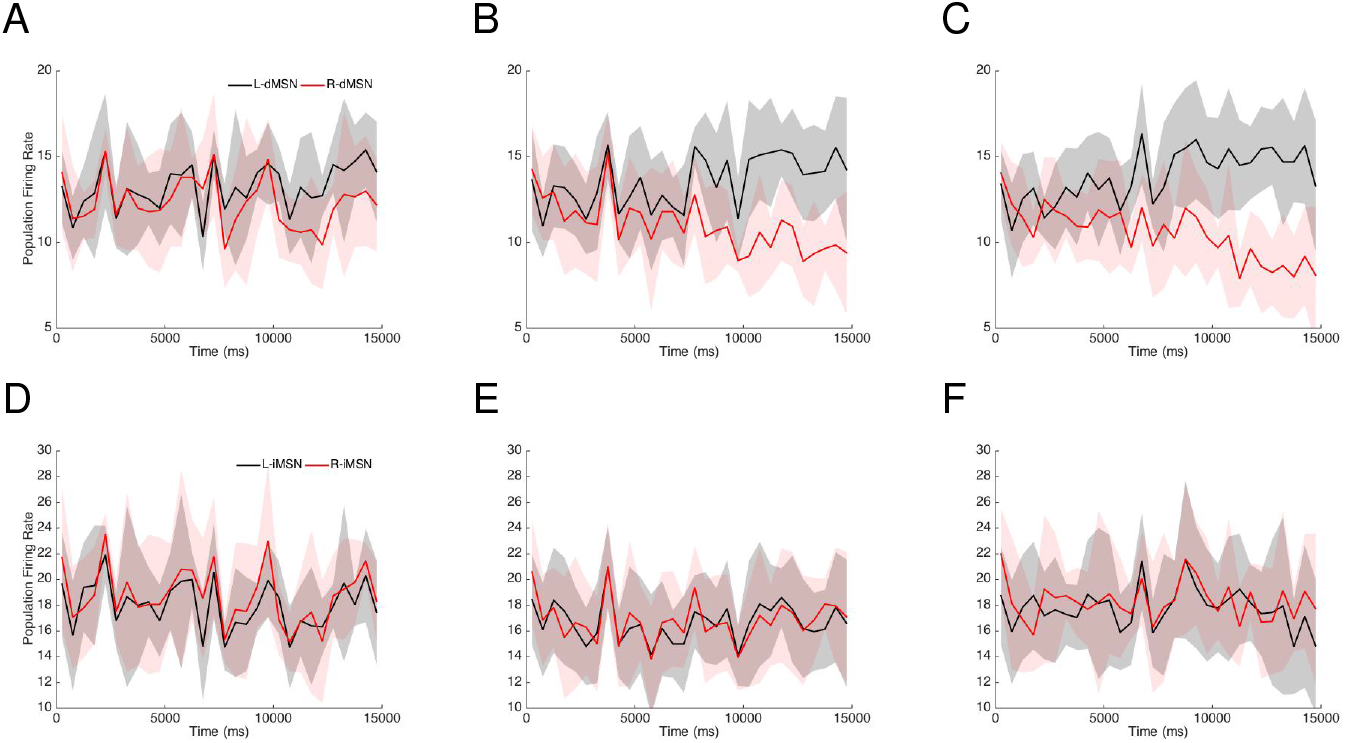
Firing rates when the reward traces are probabilistic. First column: *p_L_* = 0.65; second column: *p_L_* = 0.75; third column: *p_L_* = 0.85 case. A, B and C: Time courses of firing rates of the dMSNs selecting the *L* (black) and *R* (red) actions (50 *ms* time discretization). D, E, and F: Time courses of firing rates of the iMSNs countering the *L* (black) and *R* (red) actions (50 *ms* time discretization). In all cases, we depict the mean averaged across 8 different realizations, and the transparent regions represent standard deviations.

Similar trends across conflict levels arose in the respective frequencies of selection of action *L*. Over time, as weights to *L*-dMSN neurons grew and their firing rates increased, action *L* was selected more often, becoming gradually more frequent than action *R*. Not surprisingly, a significant difference between frequencies emerged earlier, and the magnitude of the difference became greater, for larger *p_L_* (Fig. 9).

**Fig. 9.**
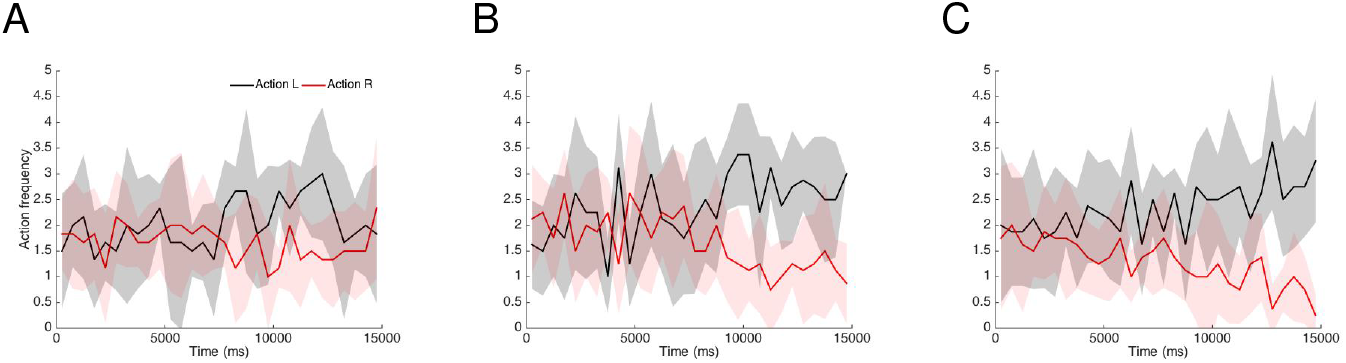
Action frequencies when reward delivery is probabilistic. All panels represent the number of *L* (black) and *R* (red) actions performed across time (discretized each 50 *ms*) when action selection is rewarded with probability *p_L_* = 0.65 (A), *p_L_* = 0.75 (B), or *p_L_* = 0.85 (C) with *p_L_* + *p_R_* = 1. Traces represent the means over 8 different realizations, while the transparent regions depict standard deviations.

To show that this feedback learning captured experimental observations, we performed additional probabilistic reward simulations to compare with behavioral data in forced-choice experiments with human subjects [20]. Each of these simulations represented an experimental subject, and each action selection was considered as the outcome of one trial performed by that subject. After each trial, a time period of 50 *ms* was imposed during which no cortical inputs were sent to striatal neurons such that no actions would be selected, and then the full simulation resumed. For these simulations, we considered the evolution of the value estimates for the two actions either separately for each subject (Fig. 10A) or averaged over all subjects experiencing the same reward probabilities (Fig. 10B2), as well as the probability of selection of action *L* averaged over subjects (Fig. 10C2).

**Fig. 10.**
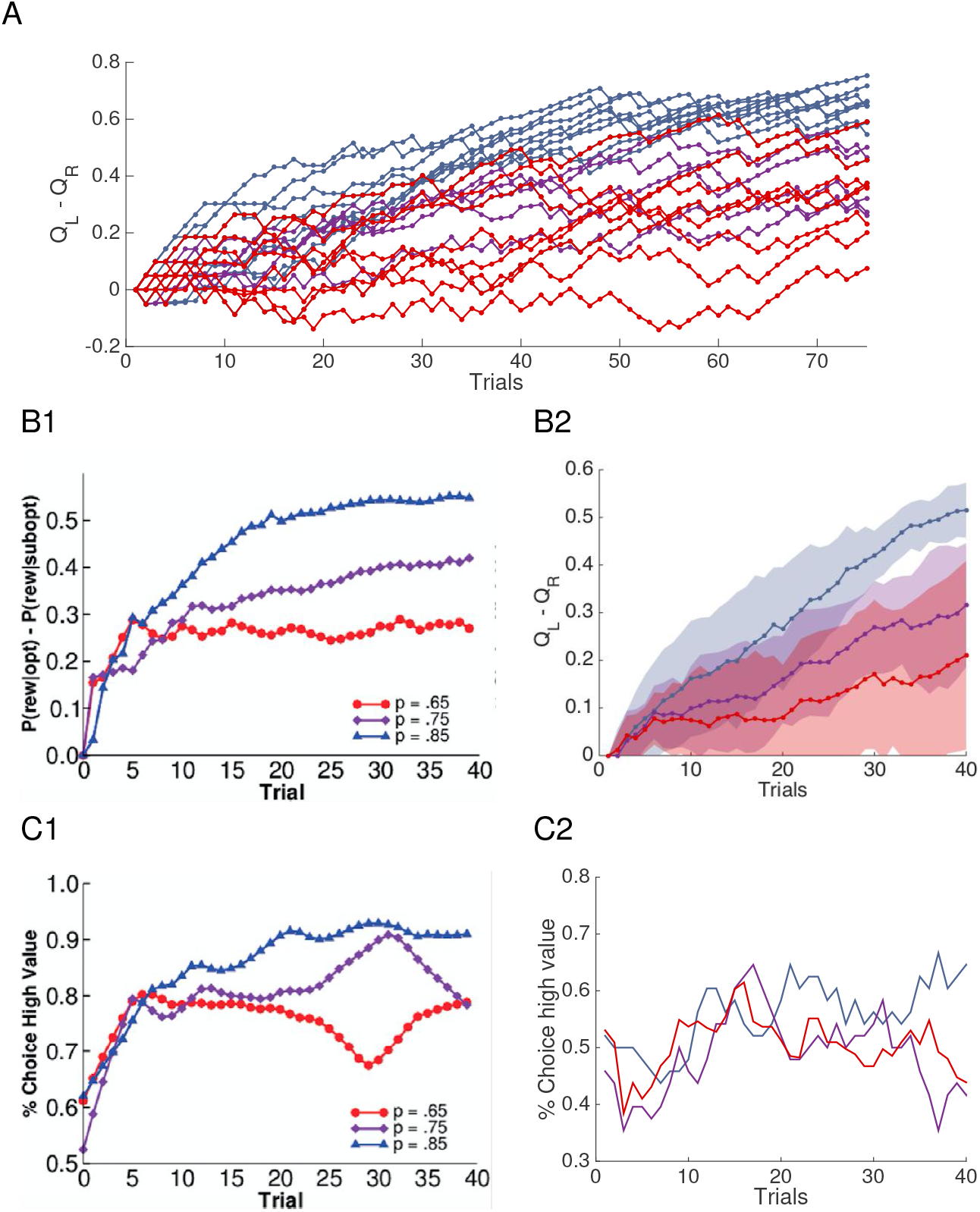
Relative action value estimates and action selection probabilities. Action value and selection probabilities were estimated over simulated trials given probabilistic reward schedules, with *p_L_* = 0.65 (dark blue), *p_L_* = 0.75 (cyan), *p_L_* = 0.85 (yellow) and *p_wL_* + *p_R_* = 1. A: Difference in action value estimates over trials in a collection of individual simulations. B: Means and standard deviations of difference in action value estimates across 8 simulations. C: Percent of trials on which the *L* action with higher reward probability was selected. B1 and C1 are the results obtained by Frank et al. in [20], obtained from 15 human subjects. B2 and C2 are the results obtained using our learning rule, combined across 8 simulations.

The mean in the difference between the action values gradually tended toward the difference between the reward probabilities for all conflict levels. Although convergence to these differences was generally incomplete over the number of trials we simulated (matched to the experiment duration), these differences were close to the actual values for many individual subjects as well as in mean (Fig. 10A,B2). These results agree quite well with the behavioral data in [20] (Fig. 10B1) obtained from 15 human subjects, as well as with observations from similar experiments with rats [50].

Also as in the behavioral experiments, the probability of selection of the more rewarded action grew across trials for all three reward probabilities, with less separation in action selection probability than in action values across different reward probability regimes (Fig. 10C2). Although our actual values for the probabilities of selection of higher value actions did not reach the levels seen experimentally (Fig. 10C1), this likely reflected the non-biological action selection rule in our STDP model (see Subsection 2.3).

## 4 Discussion

Flexible control of behavior in dynamic environments requires using the outcomes of previous actions to guide how future sensory-driven actions are chosen. In this work, we use a computational model to study the plastic effects at the corticostriatal synapses that contribute to the adaptive decision-making process. In particular, we show how a simple, dopamine-mediated STDP rule can modulate the sensitivity of both dMSN and iMSN populations to cortical inputs. This allows for the network to naturally discover which of the two targets is more likely to deliver a higher reward by modifying the ratio of direct and indirect pathway corticostriatal weights within each action channel.

Mechanistically, the adaptive selection of more rewarding actions in our simulations results from dopamine-induced increases in the efficacy of cortical synaptic inputs to dMSNs of the pathway associated with the optimal choice, while at the same time decreasing the efficacy of dMSN synapses associated with the sub-optimal choice. The changes in synapses of dMSNs coincide with a complimentary change in iMSN corticostriatal synapses, with greater decreases in weights for the optimal action channel than the sub-optimal channel. While the overall changes in iMSN weights was much smaller than the changes in dMSNs, the ratio of dMSN to iMSN weights within an action channel was a strong predictor of how the network adjusted its action selection policy to reflect learned action value estimates. These results represent testable predictions at multiple levels for how feedback learning should influence the decision process over time in the direct pathway itself. Specifically, we predict that feedback-dependent reward learning induces more salient changes in cortical synaptic weights to dMSN populations than to iMSN populations, with weights to iMSN populations remaining similar across channels.

We also characterize how plasticity driven by phasic dopamine yields rapid action value learning robustly across reward scenarios (i.e., constant rewards and probabilistic rewards). In particular, in the constant reward case, we found that when the action that elicits a higher reward (i.e., the optimal action) is switched after learning is consolidated, the network is capable of learning to favor the new optimal action, but with a longer learning time required than for the initial, unbiased learning process. In the probabilistic reward scenario, we compared our results with human experimental data obtained in previous work [20]. For different levels of conflict, or similarity of reward probabilities between the two actions, the mean in the difference between the values assigned to the actions in our model approaches the difference between the reward probabilities, matching the experimental results [20] (Figure 10A,B). On the other hand, the percentage of trials on which the higher reward action is selected is less directly related to the reward conflict level in both our simulations and the earlier experiments (Figure 10C). Overall, we find that corticostriatal learning based on reward-related dopaminergic feedback is sufficient to capture the major trends in human performance in a two-alternative forced choice task with probabilistic rewards.

The plasticity model used here makes a very compelling case for how dopamine-mediated STDP at the corticostriatal synapses can naturally modulate both firing rates and action selection in a reinforcement learning context; however, there are several improvements that can and should be the focus of future work. For example, while our model features rather detailed plasticity mechanisms that build on past modeling studies [1,22,25], we used a phenomenological action selection rule. The simplicity of the action selection mechanism is not critical to our plasticity results, since it allows higher MSN firing rates to translate into more frequent action selection. It does, however, affect the amount of time and number of trials needed for plasticity and action bias to emerge (cf. Figure 10C). We leave explicit modeling of the CBGT circuit, in the context of feedback-dependent learning and action selection, for other work [?,51]. In addition, our simulations here largely ignored the role of tonic dopamine in the action selection process [4,6,22,23,43,46], which could reflect motivation or other aspects of reward valuation in the action selection process [6,36]. Future variants of the current model should explore the influence of tonic dopamine so that we can study how both phasic and tonic dopamine mechanisms coexist and contribute to plasticity of corticostriatal connections and subsequent behavior. This addition may help to explain the role of dopamine D2 receptors in reversal learning [29]. Finally, our simulation here was limited to a very simple variant of the two-armed bandit task. While this task is a popular test of learning in the reinforcement learning literature [47], it has limited ecological validity in real world context. Future work should the dynamic range of dopamine-mediated STDP rules in more complex decision-making contexts that involve more alternatives and more complicated changes in reward dynamics.

## Acknowledgments

CV is supported by the Ministerio de Economía, Industria y Competitividad (MINECO), the Agencia Estatal de Investigation (AEI), and the European Regional Development Funds (ERDF) through projects MTM2014-54275-P, MTM2015-71509-C2-2-R and MTM2017-83568-P (AE/ERDF,EU). JR received support from NSF awards DMS 1516288, 1612913 (CRCNS), and 1724240 (CRCNS). TV received support from NSF CAREER award 1351748. The research was sponsored in part by the U.S. Army Research Laboratory, including work under Cooperative Agreement Number W911NF-10-2-0022, and the views espoused are not official policies of the U.S. Government.

